# The role of VISTA engagement in limiting neutrophil-mediated inflammation

**DOI:** 10.1101/2024.05.05.592601

**Authors:** Elizabeth C. Nowak, Jiannan Li, Mohamed A. ElTanbouly, Wilson L. Davis, Petra Sergent, Lindsay K. Mendyka, J. Louise Lines, Nicole C. Smits, Rodwell Mabaera, Shibani Rajanna, Catherine Carriere, Brent Koehn, Bruce R. Blazar, Christopher M. Burns, Randolph J. Noelle, Sladjana Skopelja-Gardner

## Abstract

A growing body of evidence suggests that VISTA, an immune checkpoint inhibitory receptor, plays a central role in the regulation of innate immunity in the settings of inflammatory diseases and cancer. Neutrophils are among the cells that have the highest membrane density of surface VISTA. Targeting VISTA on neutrophils with an agonist antibody resulted in a striking reduction in their LPS-induced peripheral accumulation. Fc receptor engagement was required for anti-VISTA antibody to mediate its effects on neutrophils. Concomitant with reduced peripheral neutrophil cell numbers, anti-VISTA antibody treatment increased neutrophil cell death in the liver. In a murine model of neutrophil-mediated arthritis, agonist anti-VISTA antibody treatment ameliorated disease severity, which was associated with reduced myeloperoxidase activity in the joints. These studies add to a growing spectrum of negative regulatory functions that VISTA performs in controlling inflammation through the innate and adaptive arms of the immune system that has implications for translation into the clinic.

## Introduction

Numerous studies have underscored the critical role for neutrophils as effectors of innate immunity (1–4). More recently, their roles as important regulators of chronic inflammation and adaptive immunity have also been elucidated (5–7). While the cascade of events regulating neutrophil recruitment and initiation of the inflammatory response has been explored at the cellular and molecular levels, the role negative checkpoint regulators play in neutrophil activation and chemotaxis remains largely unknown.

Our group has extensively studied V-domain Ig suppressor of T cell activation (VISTA (8), also known as PD-1H (9) and DD1α (10)), a negative checkpoint regulator highly expressed on myeloid cells and to a lesser degree on T lymphocytes (8, 11, 12). Multiple observations suggest that VISTA may play a unique cell-intrinsic role in the function of innate immune cells, including neutrophils. First, *Vsir* deficient (*Vsir^-/-^*) mice develop markedly attenuated collagen antibody-induced arthritis, a disease model mainly driven by neutrophils (13). Similarly, VISTA deficiency ameliorates nephrotoxic nephritis, an effect associated with reduced neutrophil activation (14). VISTA deficiency or blockade significantly reduced the presence of granulocytic myeloid derived suppressor cells (G-MDSCs) in tumors (12). In macrophages, its loss alters chemokine receptor recycling and subsequent migration (15). Finally, several reports demonstrated an important role for VISTA in MDSC-mediated suppression of T cell responses in both human and murine systems (16–18). Therefore, while accumulating evidence suggests that VISTA plays an important role in regulating innate immune cells and inflammation, if and how targeting VISTA on neutrophils shapes their response to different inflammatory stimuli has not been addressed.

In this study, we provide the first insights into a profound and primary role of VISTA in regulating neutrophil accumulation in response to systemic inflammatory stimuli. Here, we show that antibody-mediated *in vivo* targeting of VISTA decreases peripheral neutrophil levels in both LPS – and chemokine (CXCL2) – stimulated inflammation. The suppressive effects of engaging VISTA require VISTA expression on neutrophils and the expression of Fc receptor γ (FcRγ) on other cells. Anti-VISTA antibody increases neutrophil cell death, particularly in the liver. Finally, we show that antibody targeting of VISTA attenuates K/BxN serum transfer arthritis, a model dependent upon neutrophil migration to the affected joints (19). These results have broad implications for both the fundamental understanding of neutrophil persistence in peripheral tissues and therapeutic targeting of VISTA in neutrophil-mediated inflammatory diseases.

## Results

### Treatment with anti-VISTA antibody suppresses LPS-induced neutrophil accumulation in the periphery

VISTA is a negative checkpoint regulator that is expressed on both myeloid cells and T cells, with a higher expression on granulocytes and monocytes in the periphery (8, 9, 11). To assess if VISTA was expressed on neutrophils in the bone marrow, the site of their generation, or was upregulated upon their entry into the periphery, bone marrow and spleen cells were stained for VISTA expression. Robust VISTA expression was observed in neutrophils from both sites (**Figure 1A**), suggesting anti-VISTA antibodies could directly target neutrophils.

**Figure 1.**
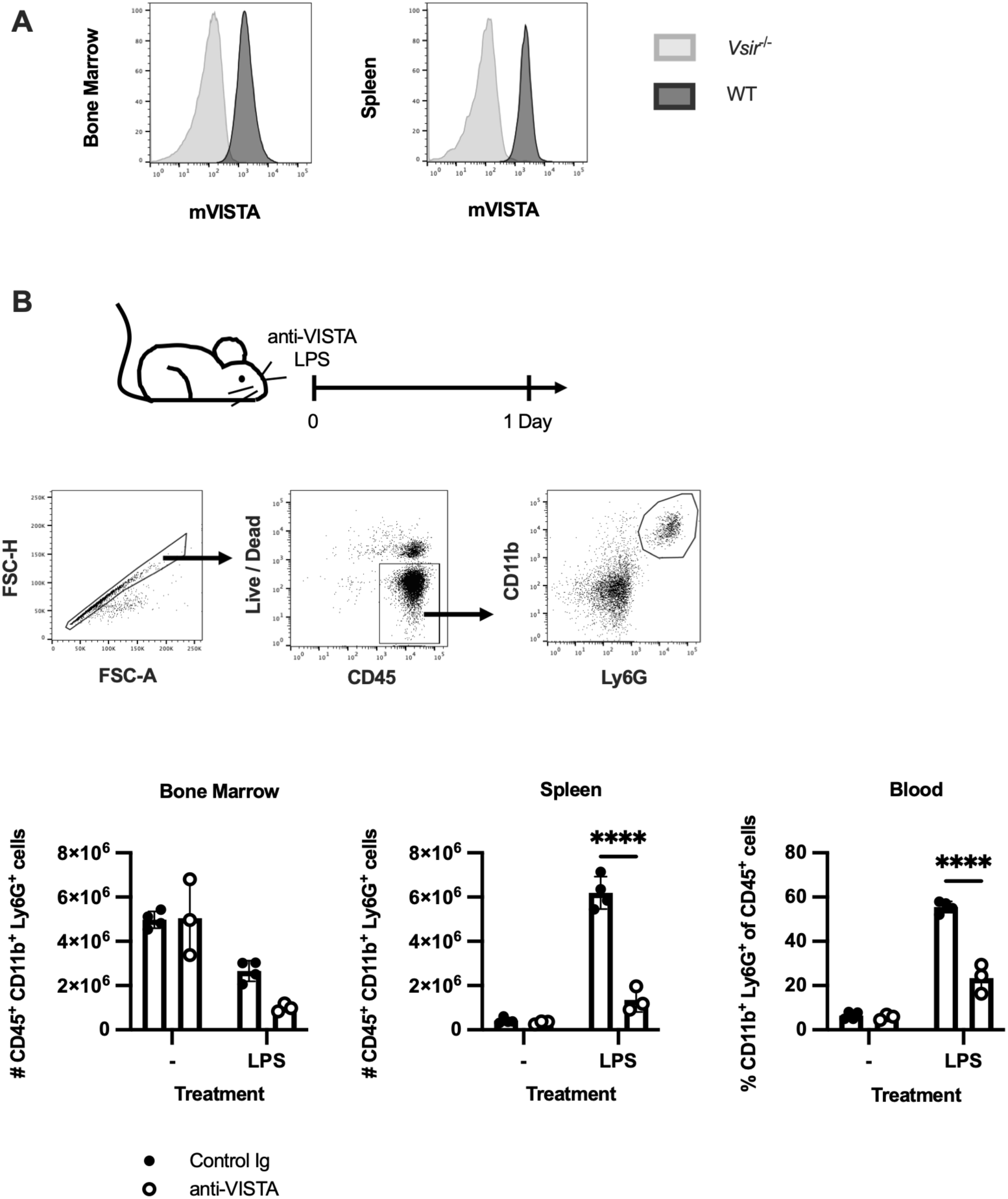
Treatment with anti-VISTA antibody suppresses LPS-induced neutrophil accumulation in the periphery. **(A)** Representative histograms show mouse VISTA (mVISTA) expression on bone marrow and spleen neutrophils (CD45^+^CD11b^+^Ly6G^+^) from WT (B6) mice (black line and dark grey shading). Control staining (light grey line and shading) is from the indicated tissue of a *Vsir*^-/-^ mouse. **(B)** B6 mice were treated i.p. with LPS (10 μg) or vehicle and anti-VISTA or control antibodies (200 μg). Dot plots show representative neutrophil gating strategy in spleen. Graphs show mean ± SD live neutrophil numbers (bone marrow and spleen) and frequency (blood) in mice 1 day after antibody treatment and LPS. Data are representative of 2 experiments: n=3-4 mice/group. Asterisks indicate statistically significant comparisons determined by 2-way ANOVA with Sidak’s multiple comparison’s test: **** p<0.0001.

To test if targeting VISTA impacts neutrophil responsiveness to an inflammatory challenge, we treated B6 mice with anti-VISTA monoclonal antibody (mAb) immediately following systemic administration of LPS (**Figure 1B**). *In vivo* stimulation with the toll-like receptor 4 ligand LPS results in the release of chemokines and cytokines, such as tumor necrosis factor, interleukin-6, CXCL2, and CXCL5, that cause activation and mobilization of neutrophils into the periphery (20–22). Mice treated with anti-VISTA mAb had significantly decreased neutrophil levels in the spleen and blood one day after LPS challenge, compared to the control group (**Figure 1B**). This was not due to the lack of neutrophil recruitment from the bone marrow, as the LPS challenge decreased neutrophil numbers in the bone marrow of both anti-VISTA and control antibody-treated mice to the similar extent (**Figure 1B**). Similar effect of targeting VISTA was subsequently observed in human VISTA knock-in (hVISTA^KI^) mice, which express human VISTA in the mouse locus of this gene (23). Anti-human VISTA (hVISTA) antibody significantly decreased neutrophil levels in the spleen and bone marrow following LPS challenge, compared to the control antibody, without suppressing neutrophil egress from the bone marrow (**Supplemental Figure 1**). Together, these data suggest that anti-VISTA antibody mediates its effects on neutrophil accumulation in the periphery, rather than on neutrophil egress from the bone marrow.

### VISTA-deficiency or anti-VISTA antibody do not impact neutrophil numbers or distribution in the absence of inflammation

The impact of VISTA deficiency and agonistic antibody targeting of VISTA on neutrophil steady-state numbers, trafficking, and distribution was investigated. No differences in neutrophil numbers (bone marrow and spleen) or frequency (blood) were observed between wild-type (B6) and *Vsir*^-/-^ (VISTA-deficient) mice at steady state (**Supplemental Figure 2**). This is consistent with a previous report showing a minor increase in granulocyte colony forming potential from VISTA deficient bone marrow in comparison to wild-type controls (15). In addition, treatment of B6 mice with anti-VISTA mAb did not affect neutrophil levels at steady state in the bone marrow, spleen, or blood one day following treatment, relative to mice treated with the control antibody (**Figure 1B**). Similar findings were observed in hVISTA^KI^ mice; treatment with anti-hVISTA mAb in the absence of an inflammatory stimulus did not change neutrophil levels in the bone marrow, blood, or spleen (**Supplemental Figure 1**). Granulocyte monocyte progenitors (defined here as cKit^+^ Sca1^-^ CD16/32^hi^ CD115^-^ CX3CR1^-^ Lineage^lo^ cells) were also not altered by antibody treatment in comparison to controls at steady state (**Supplemental Figure 3**).

### VISTA expression on neutrophils is required for anti-VISTA antibody to suppress LPS-induced neutrophil accumulation in the periphery

To determine if the effects of anti-VISTA antibody observed *in vivo* require neutrophil expression of VISTA, mice carrying neutrophil specific VISTA deletion (S100a8^cre^*Vsir*^fl/fl^) were generated. S100a8-cre expression is restricted to neutrophils and a proportion of their progenitors (24). Flow cytometry analysis confirmed the predicted loss of VISTA expression on neutrophils in the bone marrow and spleen of S100a8^cre^ *Vsir*^fl/fl^ mice, compared to cre negative (*Vsir*^fl/fl^) controls (**Figure 2A**). As expected, anti-VISTA antibody treatment following LPS challenge resulted in reduced neutrophil numbers in the spleen and blood of cre negative (*Vsir*^fl/fl^) mice (**Figure 2B**), comparable to that seen in wild-type B6 mice. In contrast, anti-VISTA mAb treatment of S100a8^cre^ *Vsir*^fl/fl^ mice following LPS stimulation showed no significant impact on neutrophil levels in the spleen and blood (**Figure 2B**). Therefore, VISTA expression by neutrophils is critical for the *in vivo* effects of anti-VISTA antibody treatment.

**Figure 2.**
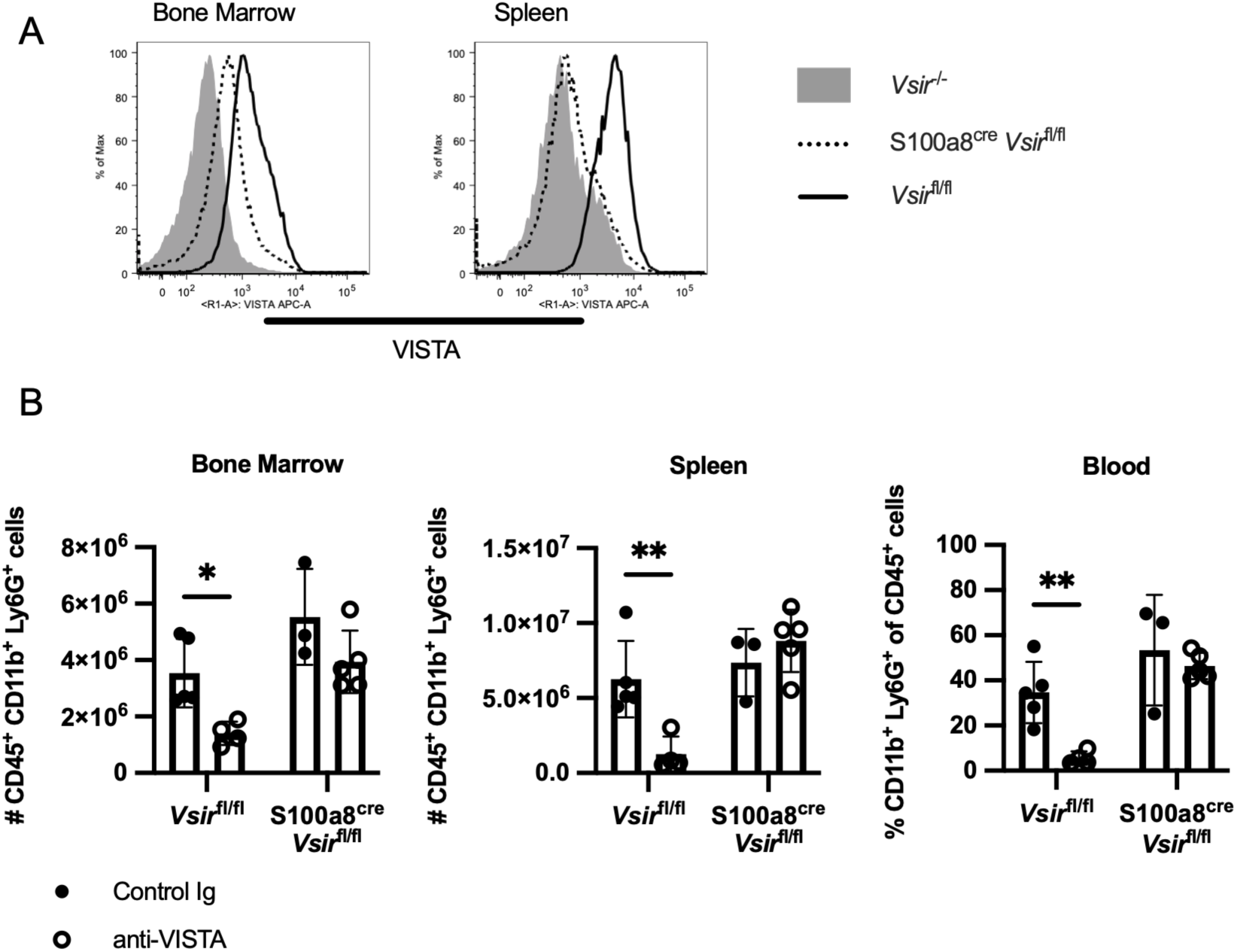
VISTA expression on neutrophils is required for anti-VISTA antibody to suppress LPS induced neutrophil accumulation in the periphery. **(A)** Histograms of VISTA expression on neutrophils (CD45^+^CD11b^+^Ly6G^+^) from *Vsir*^-/-^ (shaded area), *Vsir*^fl/fl^ (solid line), and S100a8^cre^ *Vsir*^fl/fl^ (dashed line) are shown from bone marrow and spleen. **(B)** Mice were treated with anti-VISTA or control antibody and LPS as in Figure 1 and analyzed on day 1 after treatment to determine the number (bone marrow and spleen) and frequency (blood) of live neutrophils (n=3-5 mice/group). Data are representative of 2 experiments. Graph shows mean ± SD and the result of a two-way ANOVA and Sidak’s multiple comparisons test. Asterisks indicate statistically significant comparisons: * p<0.05 and ** p<0.01.

### FcRγ engagement is required for anti-VISTA antibody to suppress LPS-induced neutrophil accumulation in the periphery

One critical consideration of antibody treatment is that the binding of the Fc portion of the antibody to the Fc receptor (FcR) can influence its mechanism of action and the downstream effects on the targeting cell. This could work through a variety of mechanisms including but not limited to crosslinking of the target antigen leading to apoptosis and / or clearance of antibody bound targets via phagocytosis or cytotoxicity (25). To initially test if FcR binding impacts the ability of anti-VISTA mAb to suppress neutrophil accumulation in response to the LPS challenge, an engineered Fc silent anti-hVISTA antibody was used in hVISTA^KI^ mice. Mice treated with the Fc silent anti-hVISTA antibody had similar neutrophil levels in the spleen following LPS stimulation as those treated with the control antibody, while mice that received the cytophilic version of anti-VISTA antibody showed a significant reduction in splenic neutrophils (**Figure 3A**). To define the contribution of specific FcRs, we evaluated the impact of anti-VISTA mAb treatment on neutrophil levels in response to LPS in *Fcer1g^-/-^* mice, which lack FcRγ and have impaired FcγRI and FcγRIII signaling (26, 27), and *Fcgr2b*^-/-^ mice, which are FcγRII deficient. We found that *Fcer1g^-/-^* mice treated with anti-VISTA mAb showed no decrease in neutrophil numbers in the spleen after LPS challenge (**Figure 3B**). In contrast, anti-VISTA mAb potently suppressed splenic neutrophil accumulation in the *Fcgr2b*^-/-^ mice (**Figure 3B**). These data suggest that FcγRI and/or FcγRIII are required for anti-VISTA mAb to prevent peripheral neutrophil accumulation in response to LPS. As neutrophils express these FcRs, we wanted to determine if FcR expression on the neutrophils themselves was necessary to cause their loss with anti-VISTA antibody treatment. Therefore wild-type and *Fcer1g^-/-^* neutrophils were isolated, labeled with different dyes, and transferred into wild-type recipient mice that were challenged with LPS and subsequently treated with anti-VISTA mAb (**Figure 3C**). We found that anti-VISTA mAb effectively suppressed splenic accumulation of both WT and *Fcer1g^-/-^* neutrophils upon transfer into wild-type hosts (**Figure 3C**), demonstrating that the lack of FcR expression on neutrophils does not impair their response to anti-VISTA mAb. Taken together, these data strongly suggest that anti-VISTA mAb coating the neutrophils binds FcR expressed by other cells to mediate neutrophil loss in the context of LPS-induced recruitment.

**Figure 3.**
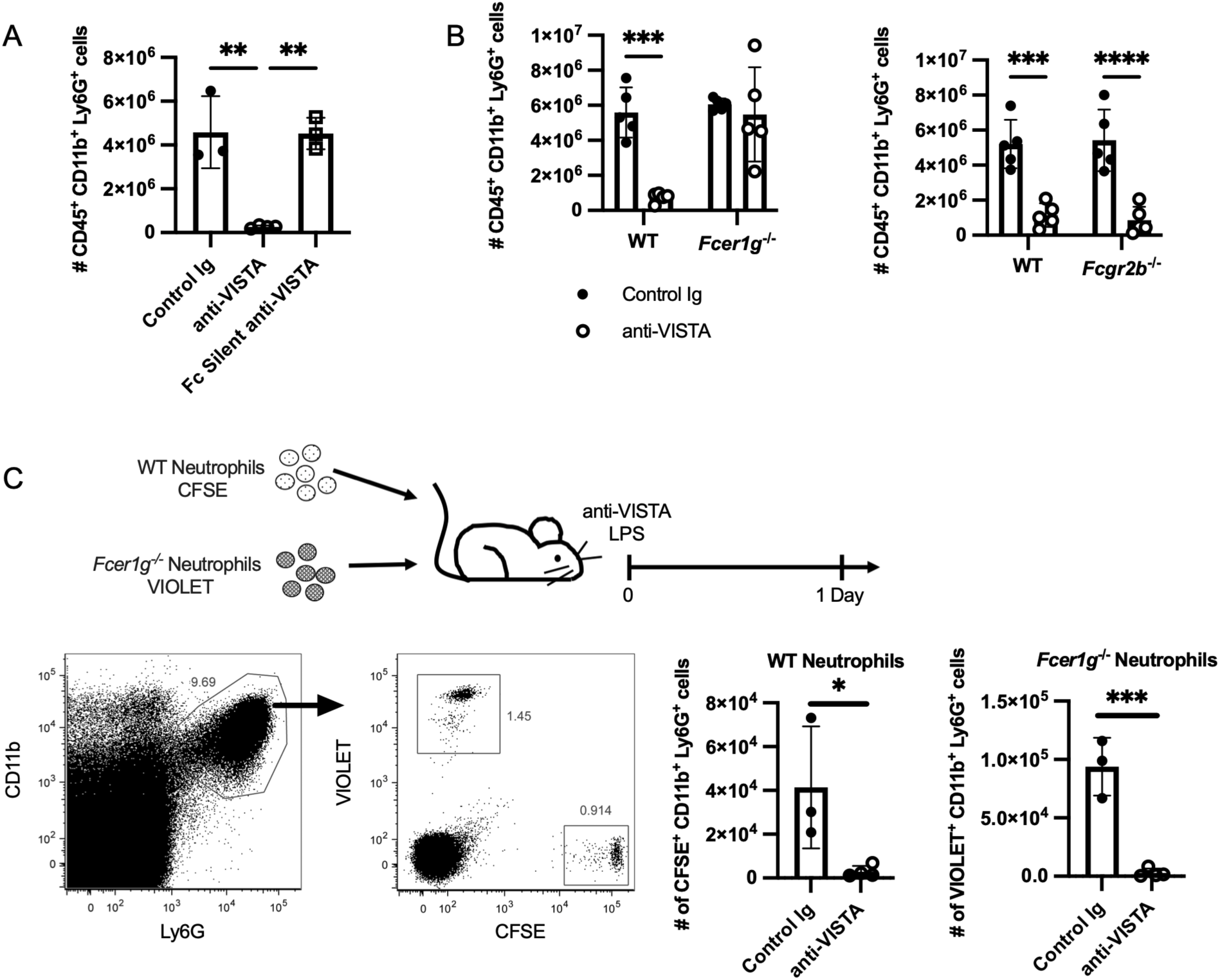
FcRγ engagement is required for anti-VISTA antibody to suppress LPS-induced neutrophil accumulation in the periphery. **(A)** Splenic neutrophil frequency was assessed 1 day post LPS treatment in human VISTA knock-in (hVISTA^KI^) mice treated with anti-VISTA or control antibodies. Data are representative of 2 independent experiments with n=3-4 mice/group. **(B)** Wild-type (B6), *Fcer1g*^-/-^, and *Fcgr2b*^-/-^ mice were treated with the anti-VISTA or control antibody and LPS as in Figure 1. Live splenic neutrophil numbers were quantified 1 day later. Data are representative of 2 independent experiments with n=5 mice/group. **(C)** Neutrophils from the bone marrow of wild-type and *Fcer1g*^-/-^ mice were isolated, labeled with CFSE or CellTrace Violet, and injected i.v. into wild-type recipients. Recipient mice were then treated with the anti-VISTA or control antibody and LPS. Plots show gating strategy from live singlet cells, and graphs show numbers of neutrophils recovered in the spleen from each transferred population. Data are representative of 3 independent experiments with n=3-4 mice/group. All graphs show mean ± SD. Statistical significance was defined by (A) one-way ANOVA with Tukey’s multiple comparisons test, (B) two-way ANOVA and Sidak’s multiple comparisons test, and (C) Student’s t-test. Asterisks indicate statistically significant comparisons: * p<0.05, ** p<0.01, *** p<0.001, and **** p<0.0001.

### Treatment with anti-VISTA antibody increases neutrophil death in the context of LPS-induced inflammation

To test if the FcR-dependent effects of anti-VISTA antibody lead to neutrophil death, bone marrow, spleen, liver, and lung were examined twelve hours post LPS challenge and antibody treatment (**Figure 4A**). The liver and lung were included as both are locations where neutrophils are recruited after LPS treatment, and the liver is a main site of neutrophil clearance (28–30). All the peripheral tissues examined had decreased live neutrophil numbers with anti-VISTA mAb treatment, indicating that the loss previously seen in the spleen was not due to the preferential trafficking to the liver or lung (**Figure 4A**). Neutrophil cell death was assessed by ApoTracker Green, a surrogate marker for Annexin V, and SYTOX, which indicates cell death through direct DNA binding. The frequency of SYTOX^+^ neutrophils was increased 2-fold in the liver with anti-VISTA mAb (**Figure 4A**). However, early apoptosis (as determined by ApoTracker Green^+^ SYTOX^-^ staining) of neutrophils in this tissue was not different between antibody treatment groups (**Supplementary Figure 4**). To test if neutrophils were forming neutrophil extracellular traps (NETs), as the increase in SYTOX+ cells would suggest, we performed immunofluorescence staining of liver for myeloperoxidase (MPO) and DNA (DAPI). MPO staining on extruded DNA strands (**Figure 4B**), confirmed the presence of NETs in the liver in the anti-VISTA mAb treatment group. Increased levels of MPO+ cells in the anti-VISTA mAb group (**Figure 4B**) coupled with a decrease in live neutrophils as detected by flow cytometry (**Figure 4A**) suggest that anti-VISTA mAb treatment promotes neutrophil clearance in the liver, potentially via NETosis.

**Figure 4.**
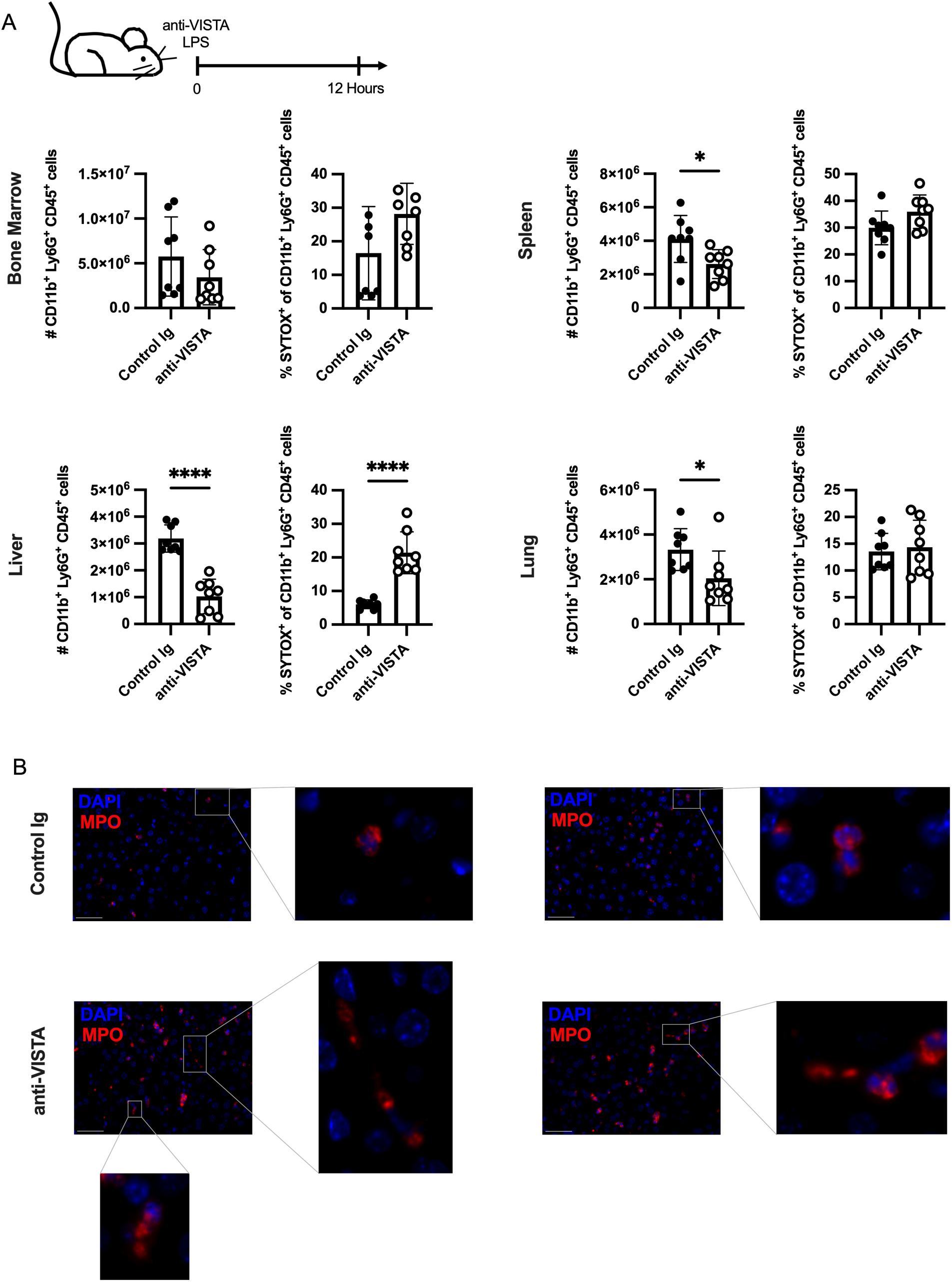
Treatment with anti-VISTA antibody in the presence of LPS increases neutrophil cell death in the liver. **(A)** B6 mice were treated i.p. with LPS (10 μg) and anti-VISTA or control antibodies (200 μg). Graphs show live neutrophil numbers and percentage of SYTOX^+^ neutrophils in the bone marrow, spleen, liver, and lung 12 hours after treatment. Data are pooled from 2 independent experiments using a total of n=8 mice/group and were analyzed by Student’s t-test. Asterisks indicate statistically significant comparisons: * p<0.05 and **** p<0.0001. **(B)** Mice were treated as described in (A) and liver tissue was processed for histology (n=4 mice/group). Tissue was stained for MPO (red) and DAPI (blue). Representative images from both groups are shown. Scale bars indicate 50 microns.

### Anti-VISTA antibody suppresses peripheral neutrophil accumulation in response to CXCL2

To ask if the effects of anti-VISTA mAb are restricted to neutrophils recruited in response to an infectious stimulus, we tested if targeting of VISTA can impact neutrophil accumulation in response to CXCL2, a critical regulator of neutrophil recruitment in response to inflammation (1, 31) that is sufficient to induce neutrophil mobilization from the bone marrow (32). *In vitro* studies demonstrated that treatment with anti-VISTA mAb inhibited the migration of both murine and human neutrophils in response to CXCL2 (**Figure 5A-B**). To confirm this *in vivo*, mice were pre-treated with anti-VISTA or control mAb and challenged with CXCL2 (**Figure 5C**). Splenic neutrophil numbers were reduced in the anti-VISTA mAb treated group one hour after CXCL2 treatment, which was paralleled by an increase in the percent of SYTOX+ neutrophils (**Figure 5C**). Even though neutrophil numbers did not differ between treatment groups in the liver, likely due to the timing of examination, anti-VISTA mAb increased neutrophil cell death, as demonstrated by a significant increase in SYTOX+ neutrophils (**Figure 5C**). These results demonstrate that anti-VISTA mAb treatment mediates cell death of neutrophils in the periphery, irrespective of the inflammatory stimulus that recruits them from the bone marrow.

**Figure 5.**
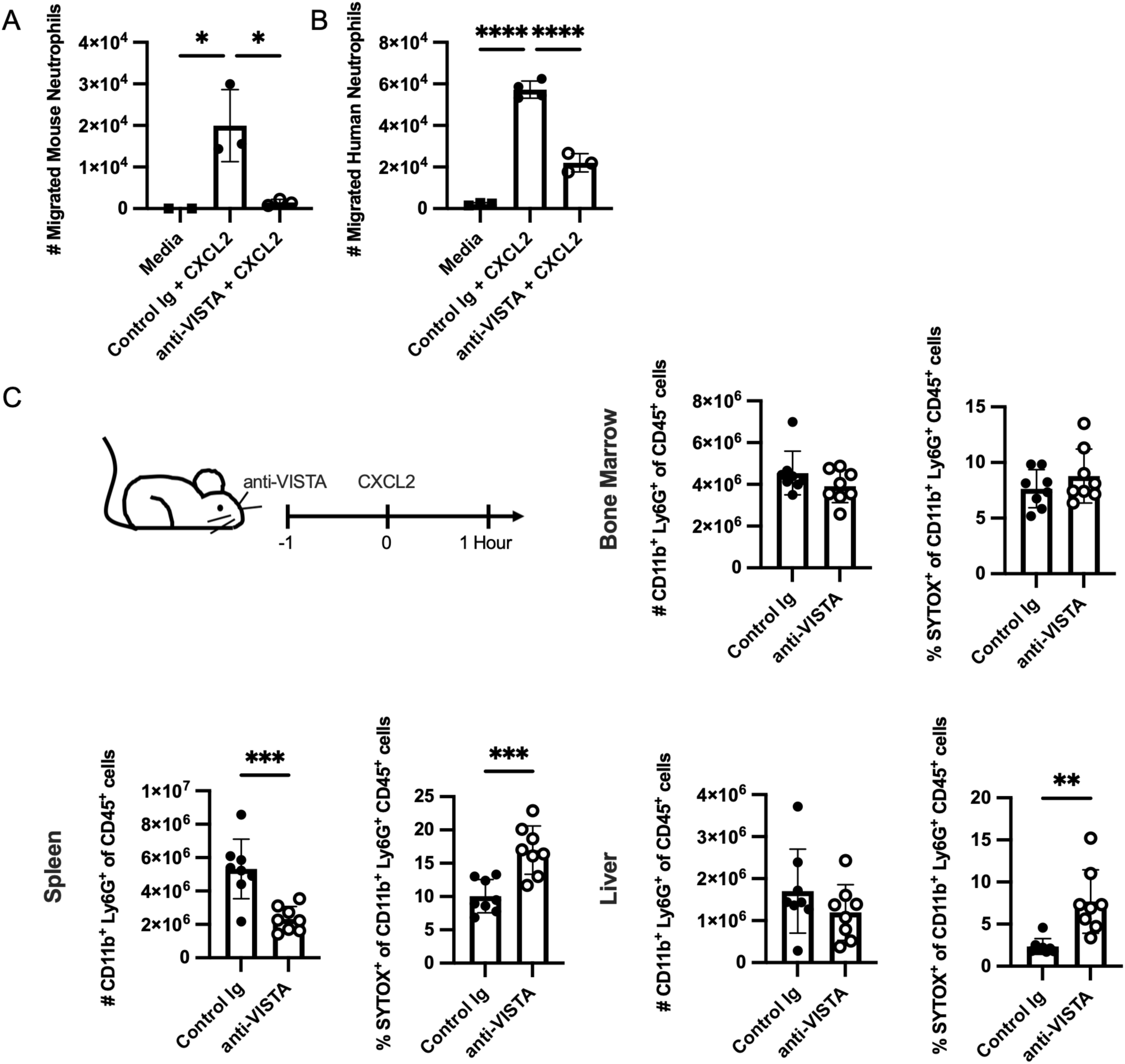
Treatment with anti-VISTA antibody reduces neutrophil response to CXCL2. **(A)** Murine neutrophils from human VISTA knock-in (hVISTA^KI^) mice were isolated from bone marrow, pre-incubated with the anti-VISTA or control antibodies, and assessed for migration to CXCL2 *in vitro*. **(B)** Human neutrophils were isolated from whole blood, pre-incubated with the indicated antibody, and assessed for migration to CXCL2 *in vitro*. Data in (A) and (B) are representative of 3 independent experiments done with 2-4 replicate wells. **(C)** Mice were treated with anti-VISTA or control antibody one hour before treatment with CXCL2 i.p. Live neutrophil numbers and percent SYTOX+ neutrophils were quantified in the indicated tissues one hour after CXCL2 injection. Data in (C) are pooled from 2 independent experiments: n=8 mice/group. Statistical significance was determined by (A, B) one-way AVONA and Tukey’s multiple comparisons test and (C) Student’s t-test. Asterisks indicate statistically significant comparisons: * p<0.05, ** p<0.01, *** p<0.001, and **** p<0.0001.

### Treatment with anti-VISTA antibody reduces the severity of K/BxN serum transfer arthritis

To examine the potential therapeutic implications of targeting VISTA to suppress neutrophil accumulation, we utilized K/BxN serum transfer arthritis. The serum donor K/BxN mice express a KRN transgenic TCR recognizing glucose-6-phosphate isomerase in the context of I-A^g7^ and develop spontaneous chronic arthritis. Transfer of serum from arthritic K/BxN mice results in transient disease in susceptible strains (33, 34). In this model, as well as in human arthritis, neutrophils play a crucial role in the induction and maintenance of disease (19, 35). We found that anti-VISTA mAb treatment markedly reduced disease severity (**Figure 6A**). To visualize neutrophil recruitment to the joints, mice were injected with luminol, a myeloperoxidase substrate that is specifically expressed in neutrophils (36). We found significant reductions in myeloperoxidase flux into the front and back paws of mice treated with the anti-VISTA mAb in comparison to control treated mice (**Figure 6B**). The flux observed in the group treated with the anti-VISTA mAb was comparable to that seen in the unimmunized (naïve) controls, indicative of the potent neutrophil elimination in the joints (**Figure 5B**). These data suggest that anti-VISTA mAb reduces disease severity by inhibiting neutrophil accumulation in the joint.

**Figure 6.**
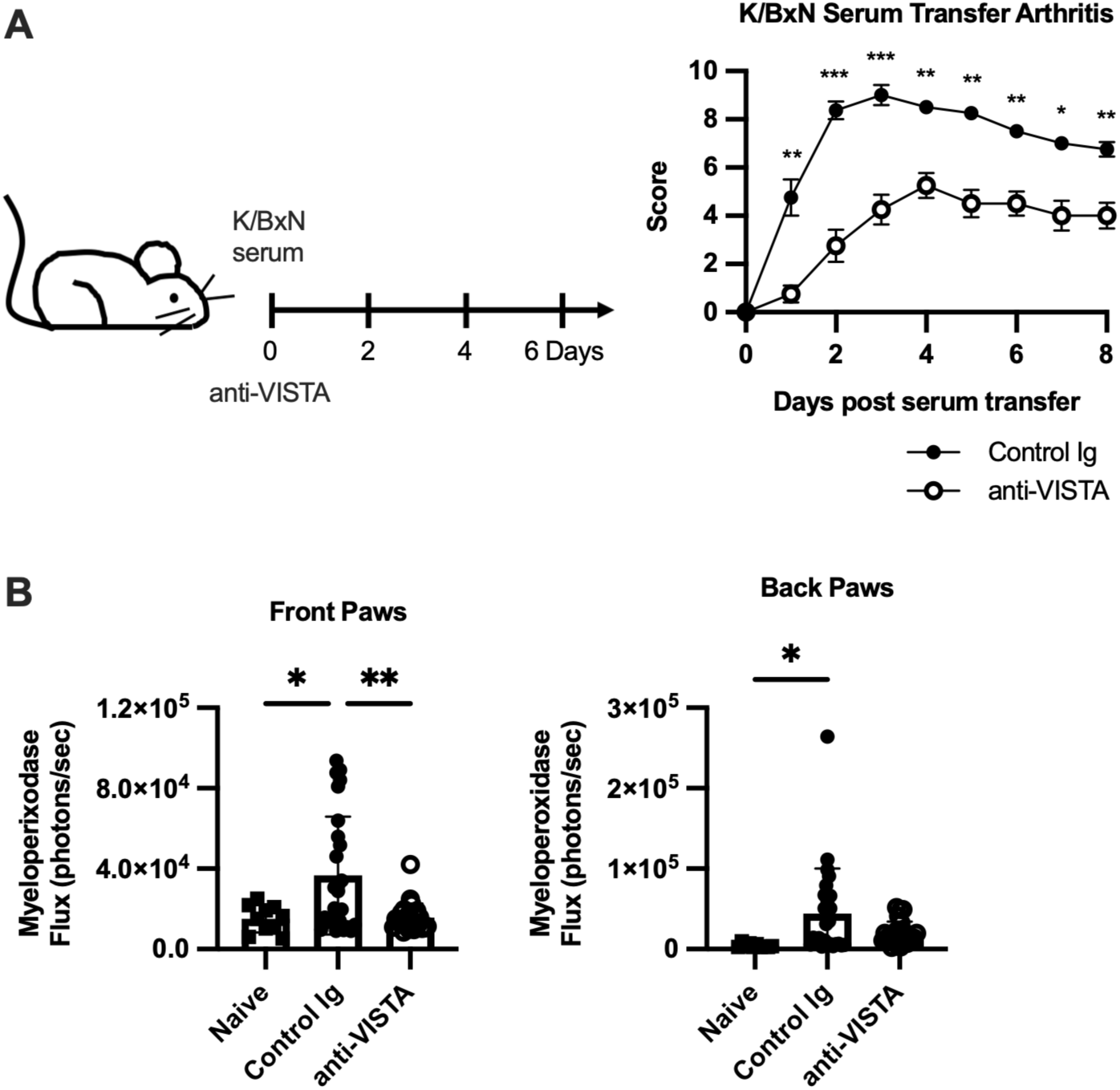
Treatment with anti-VISTA antibody reduces the severity of K/BxN serum transfer arthritis. **(A)** Mice were injected with K/BxN serum on day 0 and anti-VISTA or control antibody on days 0, 2, 4, and 6. Disease severity was monitored over time. Data are representative of 3 experiments with n=8 mice/group. Statistical significance was determined by two-way ANOVA and Sidak’s multiple comparisons test. **(B**) Mice were treated as described in (A), injected with luminol, and imaged to visualize neutrophil myeloperoxidase activity two days post serum transfer. Naïve controls were age and sex-matched mice that were not treated with K/BxN serum and antibody. Myeloperoxidase flux in indicated regions is graphed. Data were pooled from 2 independent experiments: n=5-13 mice/group. Statistics show the result of a one-way ANOVA and Tukey’s multiple comparisons test. Asterisks indicate statistically significant comparisons: * p<0.05, ** p<0.01, and *** p<0.001.

## Discussion

Here we demonstrate that agonist anti-VISTA mAb inhibits neutrophil responses by predisposing these cells to death in the peripheral tissues, following recruitment from the bone marrow. In the context of LPS stimulation, increased neutrophil cell death is observed in the liver at just twelve hours post challenge and treatment with anti-VISTA mAb. By one day post challenge the splenic neutrophil numbers are severely reduced in comparison to controls. After a challenge with CXCL2, a chemokine that mediates neutrophil egress from bone marrow in the absence of other inflammatory stimuli (32), mice given anti-VISTA mAb had decreased splenic neutrophil levels as well as increased neutrophil death in the liver and spleen.

At steady-state mature neutrophils enter the periphery due to increased CXCR2 signaling, and as they age their CXCR4 expression will increase to promote homing back to bone marrow to undergo clearance by macrophages (37). However, during inflammation bone marrow neutrophils will enter the periphery in higher numbers and these cells are recruited to the liver due to P-selectin dependent homing and eliminated via apoptosis or phagocytosis of phosphatidylserine positive neutrophils by Kupffer cells (28, 29). Under inflammatory conditions, coating neutrophils with anti-VISTA mAb may simply supply additional signals to call for their recruitment to and elimination in the liver. Future studies will elucidate if P-selectin and Kupffer cells are involved in this process. The presence of NETs, but not an increase in apoptosis, identifies NETosis as a potential mechanism by which anti-VISTA mAb leads to neutrophil death. Aspects of the mechanism of action of anti-VISTA antibody treatment on neutrophils were elucidated. An absolute requirement is direct binding of VISTA on neutrophils, which cannot be compensated for by antibody binding to other VISTA expressing cell types. Interactions with FcRγ, specifically FcγRI and FcγRIII, on cells other than neutrophils are also necessary. The implication of our findings is that antibody binding of VISTA on neutrophils allows FcR-expressing cells to mediate VISTA crosslinking, which leads to pro-apoptotic signaling cascades or to target them for phagocytosis or cytotoxicity. As previously reviewed, this is observed with some antibodies used in cancer immunotherapy (25, 38). Our studies suggest that crosslinking of anti-VISTA mAb via FcR-expressing cells leads to neutrophil elimination in the liver possibly via NETosis. The ability of antibody crosslinking to induce NETs has previously been reported and could have implications for neutrophil functions in infection (39). Future studies will address if VISTA targeting on neutrophils potentiates pathogen elimination and probe the pertinent mechanisms.

Targeting VISTA on myeloid cells with therapeutic antibodies is of great interest and promise in both cancer and inflammatory diseases (12, 13, 40). In terms of the latter conditions, treatment with agonistic antibodies may be an effective strategy to abolish the acute influx of neutrophils into inflammatory microenvironments. This approach could potentially have profound clinical benefits on diseases such as inflammatory arthritis, chronic obstructive pulmonary disease, reperfusion injury, and myocardial infarction where exuberant neutrophil responses may result in organ damage, morbidity, and in some cases mortality. In the context of cancer, eliminating neutrophils by targeting VISTA may be beneficial in suppressing neutrophil-mediated angiogenesis in the tumor environment (41).

## Materials and Methods

### Mice

*Vsir^-/-^* mice were obtained from the Mutant Mouse Resource & Research Center and bred onto a C57BL/6N (B6) background. Controls for them were C57BL/6 (Strain Code: 027) mice from Charles River Laboratories (Wilmington, MA). *Fcer1g*^-/-^ (Strain: 583), *Fcgr2b*^-/-^ (Strain: 580), and control C57BL/6 (Strain: B6) were purchased from Taconic Biosciences. *Vsir*^fl/fl^ mice were previously described and provided by Sam Lee (10). These were intercrossed with S100a8^cre^ (JAX Stock No: 021614) mice to generate S100a8^cre^ VISTA^fl/fl^ mice and cre-negative littermates. Balb/cJ (Stock No: 000651) were purchased from the Jackson Laboratory (Bar Harbor, ME). Human VISTA knock-in (hVISTA^KI^) mice were previously described (23). KRN T cell receptor transgenic mice were provided by Diane Mathis and Christopher Benoist (Harvard Medical School). They were interbred with NOD/ShiLtJ (JAX Stock No: 001976) to generate K/BxN mice. All mice were at least 8 weeks old at the start of experiment. Controls were age and sex matched across groups. Studies were approved by Dartmouth’s Institutional Animal Care and Use Committee. All animals were maintained in a specific pathogen-free facility at Geisel School of Medicine at Dartmouth.

### In Vivo Neutrophil Activation and Migration

Mice were injected i.v. with 10 μg LPS purified from *E. coli* O55:B5 (Millipore Sigma, St. Louis, MO) and treated i.p. with 200 μg Hamster Ig (Control Ig; BioXcell, Lebanon, NH), anti-mVISTA (8G8), human IgG2 (Control Ig, CB2, Crown Bioscience, Cambridge, MA), or anti-hVISTA (803), respectively. Antibodies against mouse and human VISTA were generated as previously described (8, 11). For one set of experiments an additional agonist, anti-hVISTA clone (901), and an FcR silent version of it were utilized. In adoptive transfer experiments, bone marrow was harvested from donor mice and enriched for neutrophils using a Neutrophil Isolation Kit (Miltenyi Biotec). Isolated cells were labeled with Cell Trace CFSE or Violet Proliferation Kit (ThermoFisher Scientific) according to manufacturer’s directions and injected at 5×10^5^ – 1×10^6^ cells/mouse i.v. before LPS and antibody treatment. Tissues were harvested at 12 or 20 hours after treatment. To study neutrophil migration, mice were injected with 200 μg antibody i.v. 1 hour before receiving an i.p. injection of 1 μg CXCL2 (Peprotech, Rocky Hill, NJ). Readout was 1 hour after the CXCL2 injection.

### Cell Isolation and Flow Cytometry

Harvested tissues were initially placed in Hank’s balanced salt solution. Spleens were manually dissociated, filtered, and red blood cells lysed. Bone marrow was isolated using centrifugation as described by Amend et al. (42) and underwent red blood cell lysis and filtering. Lungs were processed using a Lung Dissociation kit, mouse (Miltenyi Biotec, Bergish Gladbach, Germany), according to manufacturer’s directions. Liver was processed as described by Daemen at el. (43) except the perfusion was done manually. The cell numbers were normalized to the total weight of the liver in experiments where a portion of the liver was taken for histology and the remaining was processed to obtain cells for flow cytometry.

The following antibodies were purchased conjugated to FITC, PE, PerCP, PE-Cy7, APC, Alexa Fluor 647, APC-Cy7, Brilliant Violet 421, eFluor 450, and/or Brilliant Violet 510: CD45 (30-F11), CD11b (M1/70), Ly6G (1A8), Ly6C (HK1.4), mVISTA (MIH63), hVISTA (MIH65), c-Kit (2B8), Sca-1 (D7), CD16/32 (93), CD115 (AFS98), CX3CR1 (SA011F11), TER-119 (TER-119), CD3 (145-2C11), B220 (RA3-6B2), and CD11c (N418) from Biolegend (San Diego, CA), ThermoFisher Scientific (Waltham, MA), or BD Biosciences (Franklin Lakes, NJ).

Except for the experiments using SYTOX, cells were stained using LIVE/DEAD Fixable Near-IR or Yellow Dead Cell Stain Kit (ThermoFisher Scientific) according to manufacturer’s directions. Samples were incubated with TruStain FcX PLUS (S170011E; Biolegend) and stained with antibody cocktail. Flow cytometry was performed on MACSQuant or Accuri instruments. Data analysis was done using FlowJo software (Treestar Incorporated, Ashland, OR) following initial gating on Live/Dead negative singlets. In experiments using SYTOX Blue (ThermoFisher Scientific), cells were stained for surface markers as above, ApoTracker Green (Biolegend) according to manufacturer’s directions and SYTOX Blue was added shortly before performing flow cytometry.

### Histology

Liver tissue was fixed in 10% neutral buffered formalin and paraffin-embedded into cassettes that were used to make slides with 4 micron sections. Staining was done following the guidelines in the Opal 7-color Manual IHC Kit (Akoya Biosciences, Catalog: NEL811001KT). Briefly, slides were deparaffinized, rehydrated, fixed, and underwent antigen retrieval in AR6 buffer (via heating in a microwave). After a cool down, slides were blocked with Akoya antibody diluent/block and stained with a primary antibody against myeloperoxidase (Thermo Fisher Scientific, Catalog: PA5-16672), washed, stained with goat anti-rabbit HRP (Thermo Fischer Scientific, Catalog: 31462), washed, incubated with Opal 650, washed, and underwent microwave treatment in AR6, incubated with Spectral DAPI, washed, and mounted with Prolong Diamond (Thermo Fisher, Catalog: P36961). A Vectra 3.0 microscope was used for image acquisition.

Briefly, whole slide scans were done, Phenochart was used to select MSI regions, and images were acquired at 40x magnification. Spectral libraries were generated using single color control slides for Opal 650 and DAPI and images were unmixed using InForm 2.60 software. ImageJ software was used to add scale bars and adjust brightness and contrast.

### Human Blood Samples

Human blood samples from healthy adult volunteers were collected after signed consent in accordance with IRB–reviewed protocols. The IRB approval (Reference number D10083) was granted by Trustees of Dartmouth College Committee for the Protection of Human Subjects. The healthy subjects were all female and between the ages of 24-68 years at the time of blood-draw. None of the donors had a documented history of SARS-CoV or SARS-CoV-2 infection.

### In Vitro Neutrophil Migration

Bone marrow was harvested from B6 mice and neutrophils purified using Neutrophil Isolation Kit (Miltenyi Biotec) according to manufacturer’s directions. Cells were incubated with 30 μg/ml of indicated antibody for 1.5 hours at 37°C. Transwell plates (24 wells with 3 μm pore membrane; Corning, Corning, NY) were loaded with 20 ng CXCL2 in the bottom compartment and 6×10^5^ neutrophils in the top compartment. Plates were incubated at 37°C for 45 minutes before neutrophils in the bottom chamber were quantified by FACS.

For human neutrophil migration, cells were purified from blood using an EasySep Human Neutrophil Isolation Kit (Stemcell Technologies, Vancouver, Canada) according to manufacturer’s directions. Isolated cells were incubated with 10 μg/ml of indicated antibody for 4 hours at 37°C.

Transwell plates were loaded with 120 ng CXCL2 in the bottom compartment and 4×10^5^ neutrophils in the top compartment. Plates were incubated at 37°C for 1 hour before cells from the bottom chamber were quantified by FACS.

### K/BxN Serum Transfer Arthritis

Serum from donor K/BxN mice was isolated after mice had developed spontaneous arthritis. To induce arthritis in Balb/cJ mice, pooled K/BxN serum was injected into mice at 200μl/mouse i.p. on day 0. Each paw was scored every day over 8 days as follows: 1 – swelling in 1 joint type (considering ankle, foot, and toes), 2 – swelling in 2 joints, and 3 – swelling in all 3 joints. Total score for a mouse was the sum of score of all 4 paws. For these experiments, mice were treated with 300 μg/mouse with indicated antibody. In some experiments, mice were given luminol at 100 mg/kg and imaged on a Xenogen IVIS machine to assess flux (photons/second).

## Author Contributions

R.J.N., S.S.G., C.M.B., and E.C.N. designed the research studies. E.C.N., J.L., M.A.E., W.L.D., P.S., L.K.M., J.L.L., N.C.S., R.M., S.R., C.C., and B.K. conducted experiments and/or analyzed data. E.C.N., S.S.G., and R.J.N. wrote the manuscript, and C.M.B. and B.R.B. critically reviewed and edited it.

## Acknowledgements

This work was performed with the assistance of the Immune Monitoring and Flow Cytometry Resource, Irradiation, Pre-clinical Imaging and Microscopy Resource, and the Pathology Shared Resource. These core facilities are supported by NCI Cancer Center Support Grant 5P30CA023108-37 and 5P30CA023108-43.

This work was supported by NIH grants R01CA214062 and R01AR070760 to R.J.N. and R21AR079661-01, 2022 LRA Lupus Innovation Award, and CDMRP Lupus Research Program W81XWH2110889 for S.S.G.

---

**Supplemental Figure 1.**
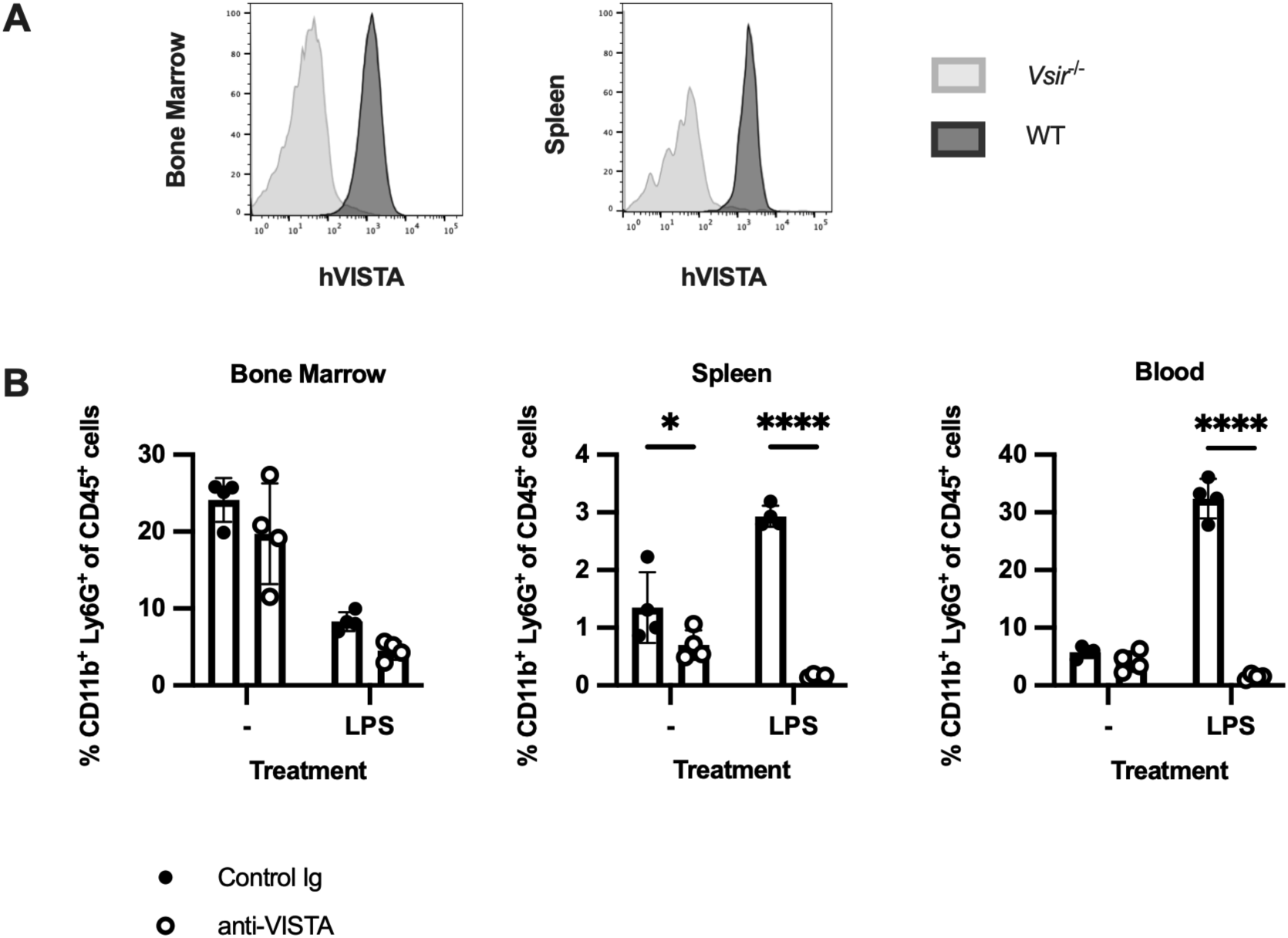
Anti-hVISTA antibody treatment given with LPS impairs neutrophil accumulation in the spleen and blood in hVISTA^KI^ mice. **(A)** Representative human VISTA (hVISTA) staining of live neutrophils from bone marrow and spleen of human VISTA knock-in (hVISTA^KI^) mice. (**B)** Mice expressing hVISTA were treated with control or anti-hVISTA antibody and LPS and analyzed on day 1 after treatment to determine neutrophil frequency in the bone marrow, spleen, and blood (n=3-5 mice/group). Data are representative of 2 experiments. All graphs show mean ± SD and the results of a two-way ANOVA and Sidak’s multiple comparisons test. Asterisks indicate statistically significant comparisons: * p<0.05 and **** p<0.0001.

**Supplemental Figure 2.**
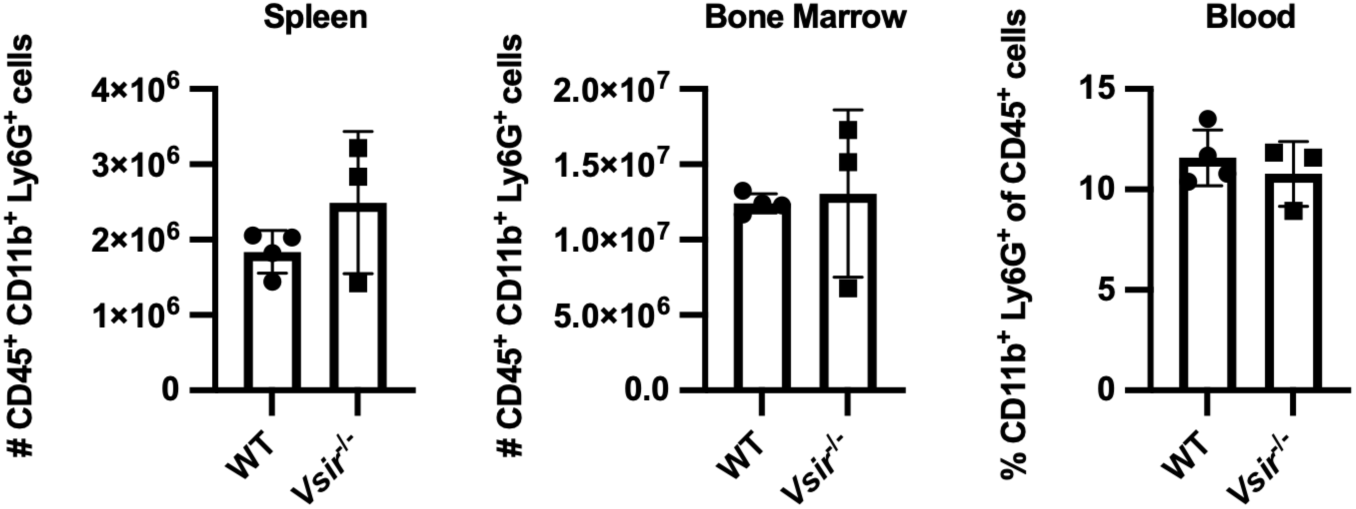
VISTA deficiency has minimal effects on neutrophil levels at steady state. Neutrophil numbers in bone marrow and spleen and neutrophil frequency in blood of wild-type (B6) and *Vsir*^-/-^ mice were quantified by flow cytometry. Results were not statistically significant by Student’s t-test (n=3-4 mice/group).

**Supplemental Figure 3.**
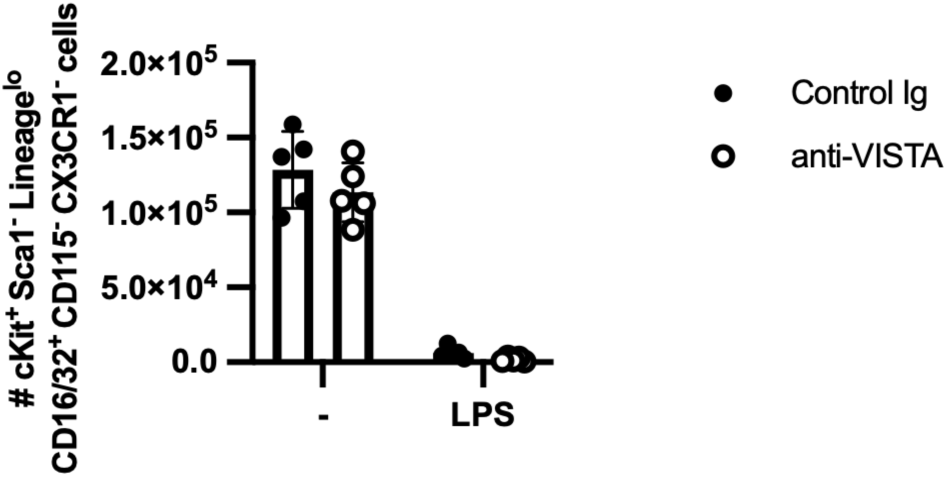
An-VISTA antibody treatment has minimal effects on granulocyte monocyte progenitors. WT mice were treated with control or anti-VISTA antibody and LPS and analyzed on day 1 after treatment to determine progenitor numbers in bone marrow. Results were not statistically significant by two-way ANOVA and Sidak’s multiple comparisons test (n=5 mice/group).

**Supplemental Figure 4.**
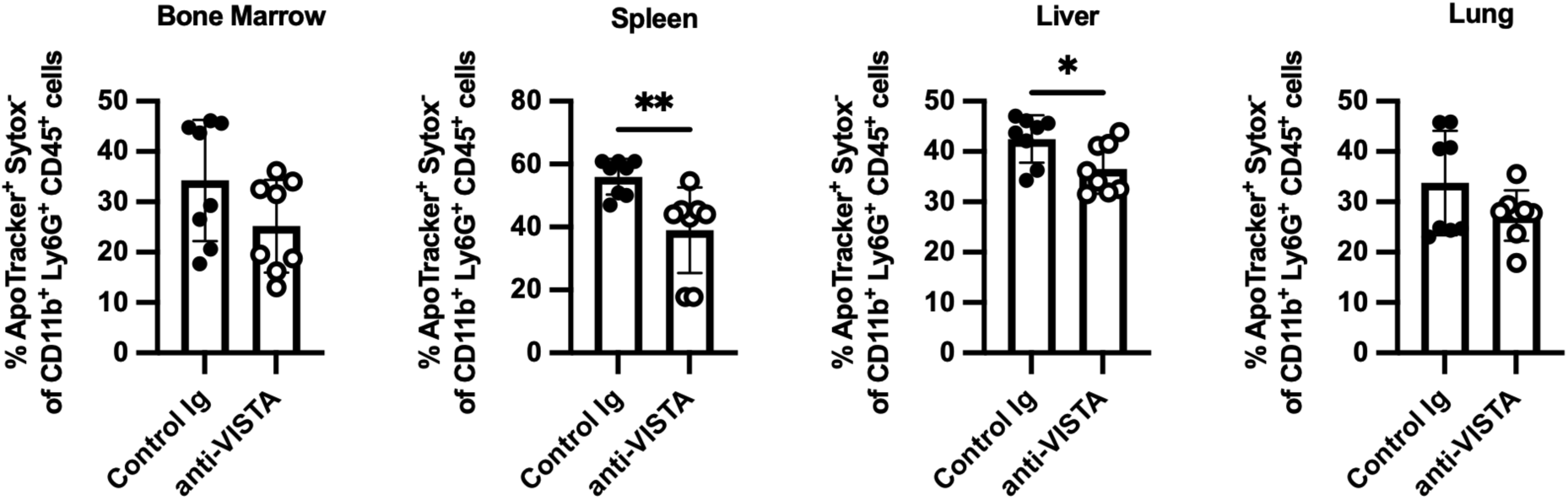
Treatment with anti-VISTA antibody in the presence of LPS does not increase early apoptosis. Graphs show the percentage of ApoTracker^+^ SYTOX^-^ cells in the indicated tissues in WT mice 12 hours after treatment with LPS and anti-VISTA or control antibody. Data are pooled from 2 independent experiments: n=8 mice/group. Asterisks indicate statistically significant comparisons by Student’s t-test: * p<0.05 and ** p<0.01.

